# *Microbe-seq*: high-throughput, single-microbe genomics with strain resolution, applied to a human gut microbiome

**DOI:** 10.1101/2020.12.14.422699

**Authors:** Wenshan Zheng, Shijie Zhao, Yehang Yin, Huidan Zhang, David M. Needham, Ethan D. Evans, Chengzhen L. Dai, Peter J. Lu, Eric J. Alm, David A. Weitz

## Abstract

We present *Microbe-seq*, a high-throughput single-microbe method that yields strain-resolved genomes from complex microbial communities. We encapsulate individual microbes into droplets with microfluidics and liberate their DNA, which we amplify, tag with droplet-specific barcodes, and sequence. We use *Microbe-seq* to explore the human gut microbiome; we collect stool samples from a single individual, sequence over 20,000 microbes, and reconstruct nearly-complete genomes of almost 100 bacterial species, including several with multiple subspecies strains. We use these genomes to probe genomic signatures of microbial interactions: we reconstruct the horizontal gene transfer (HGT) network within the individual and observe far greater exchange within the same bacterial phylum than between different phyla. We probe bacteria-virus interactions; unexpectedly, we identify a significant *in vivo* association between crAssphage, an abundant bacteriophage, and a single strain of *Bacteroides vulgatus. Microbe-seq* contributes high-throughput culture-free capabilities to investigate genomic blueprints of complex microbial communities with single-microbe resolution.

## Introduction

Microbial communities inhabit many natural ecosystems, including the ocean, soil, and in the digestive tracts of animals (Human Microbiome Project, 2012b; Lloyd-Price et al., 2017; Sunagawa et al., 2015; Thompson et al., 2017). One such complex and important community is the human gut microbiome. Comprising trillions of microbes in the gastrointestinal tract (Sender et al., 2016), this microbiome has substantial associations with human health and diseases, including metabolic syndromes, cognitive disorders, and autoimmune diseases (Cho and Blaser, 2012; Shreiner et al., 2015). The behavior and biological effects of a microbial community depend not only on its composition (Vanessa K. Ridaura et al., 2013; Yatsunenko et al., 2012), but also on the biochemical processes that occur within each microbe, and on the interplays between them (Coyte et al., 2015; Rakoff-Nahoum et al., 2016); these processes are strongly impacted by the genomes of each individual microbe living in that community.

The composition of the gut microbiome is specific to each individual; while people in many cases carry similar sets of microbial species, different individuals have distinct subspecies strains, which exhibit significant genomic differences, including point mutations and structural variations (Lloyd-Price et al., 2017; Poyet et al., 2019; Truong et al., 2017; Zhao et al., 2020). These genomic variations between strains can lead to differences in important traits, including antibiotic resistance and metabolic capabilities (Scholz et al., 2016), which can have significant consequences to human health. For example, *Escherichia coli* are common in healthy human gut microbiomes, yet particular *E. coli* strains have been responsible for several lethal foodborne outbreaks (Nataro and Kaper, 1998). Microbial behavior in the gut microbiome is influenced not only by the presence of particular strains, but also by the interactions among them, such as cooperation and competition for food sources (Rakoff-Nahoum et al., 2016), phage modulation of bacterial composition (Manrique et al., 2017; Minot et al., 2013), and transfer of genomic materials between individual microbial cells (Huddleston, 2014; Smillie et al., 2011). Improving our fundamental understanding of these behaviors depends on detailed knowledge of the genes and pathways specific to particular microbes (Human Microbiome Project, 2012a); elucidating this information can present significant challenges where taxa are only known to the species level, obscuring strain-level differences. Individual microbes from the same subspecies strain from a single microbiome largely share the same genome (Poyet et al., 2019; Zhao et al., 2019); therefore, a fundamental improvement in understanding would be provided by high-quality genomes resolved to the level of subspecies strains from a broad range of microbial taxa within the community.

Several approaches are used to explore the genomics of the human gut microbiome. One widely-used general technique is shotgun metagenomics, in which a large number of microbes are lysed and the DNA sequenced to yield a broad survey of the genomic content from the microbial community (Jovel et al., 2016; Scholz et al., 2016; Xie et al., 2016). Metagenomics-derived sequences have been assigned to individual species and have been used to construct genomes; however, metagenomics is generally unable to assign DNA sequences that are common to multiple taxa in a single sample, such as when one species has multiple strains, or when homologous sequences occur in the genomes of multiple taxa (Bishara et al., 2018; Brito and Alm, 2016; Van Rossum et al., 2020). Consequently, metagenomics cannot in general resolve genomes with subspecies strain resolution. By contrast, high-quality strain-resolved genomes of taxa from the human gut microbiome have been assembled from colonies cultured from individual microbes (Almeida et al., 2019; Browne et al., 2016; Poyet et al., 2019; Zhao et al., 2019); however, culturing colonies can be labor-intensive and biased toward easy-to-culture microbes. Alternatively, single-cell genomics relies upon isolating and lysing individual microbes in wells on a titer plate, and subsequently amplifying their DNA for sequencing (Chijiiwa et al., 2020; Lasken, 2007; Pachiadaki et al., 2019; Rinke et al., 2014; Xu et al., 2016); such approaches also yield strain-resolved genomes and have been used to probe the association between phages (DŽunková et al., 2019) and bacteria in the human gut microbiome. For both culture and well-plate approaches, however, available resources severely limit the number of strain-resolved genomes that originate from the same community (Almeida et al., 2019; Poyet et al., 2019), thereby constraining our knowledge of the genomic structure and dynamics of the human gut microbiome of an individual. Thus, substantial improvement of this understanding requires a practical, cost-effective method to obtain single-microbe genomic information at the level of detail given by culture-based or single-cell genomics, while at the same time sampling the broad spectrum of microbes typically accessed by shotgun metagenomics.

Here we introduce *Microbe-seq*, a high-throughput method to obtain the genomes of large numbers of individual microbes. We use microfluidic devices to encapsulate individual microbes into droplets, and we lyse, amplify whole-genomes, and barcode the DNA within these droplets. Consequently, we achieve substantially higher throughput than practically accessible with titer plates. We investigate the human gut microbiome, analyzing seven longitudinal stool samples collected from one healthy human subject, and acquire 21,914 single amplified genomes (SAGs). We compare with metagenomes from the same samples and find that these SAGs capture a similar level of diversity. We group SAGs from the same species and co-assemble to obtain the genomes of 76 species; 52 of these genomes are of high quality, with more than 90% completeness and less than 5% contamination. We achieve single-strain resolution and observe that six of these species have multiple strains, whose genomes we assemble. Using *Microbe-seq*, we can probe genomic signatures of microbial interactions within the community. For instance, we construct the network of the horizontal gene transfer (HGT) of the bacterial strains in a single individual’s gut microbiome and find significantly greater transfer between strains within the same bacterial phylum, relative to those in different phyla. Unexpectedly, we detect association between phages and bacteria using *Microbe-seq*: we find that the most common bacteriophage in the human gut microbiome, crAssphage (Guerin et al., 2018), has significant *in vivo* association with only a single strain of *Bacteroides vulgatus*.

## Results

### High-throughput sample preparation using droplet-based microfluidic devices

We use a microfluidic device to encapsulate individual microbes into droplets containing lysis reagents, as shown in the schematic in Figure 1A and images in Figure 1B. We collect the droplets in a tube and incubate to lyse the microbes; the DNA from each individual microbe remains within its own single droplet. We reinject each droplet into a second microfluidic device (Ahn et al., 2006) that uses an electric field to merge it with a second droplet containing amplification reagents (Blanco et al., 1989; Dean et al., 2001); we collect the resulting larger droplets and incubate them to amplify the DNA. We then use similar procedures with a third microfluidic device to merge each droplet with another droplet containing reagents to fragment and add adapters (Nextera) to the DNA (Adey et al., 2010). We subsequently employ a fourth microfluidic device to merge each droplet with an additional droplet containing a barcoding bead, a hydrogel microsphere with DNA barcode primers attached, as shown in Figure 1C; these primers contain two parts, one barcode sequence unique to each droplet, and another sequence that anneals to the previously-added adapters. We attach these barcode primers to the fragmented DNA molecules within each droplet using polymerase chain reaction (PCR). We then break the droplets, add sequencing adapters, and sequence (Illumina). We illustrate all of these steps in the schematic in Figure 1A and include schematics for all microfluidic devices (Figure S1).

**Figure 1.**
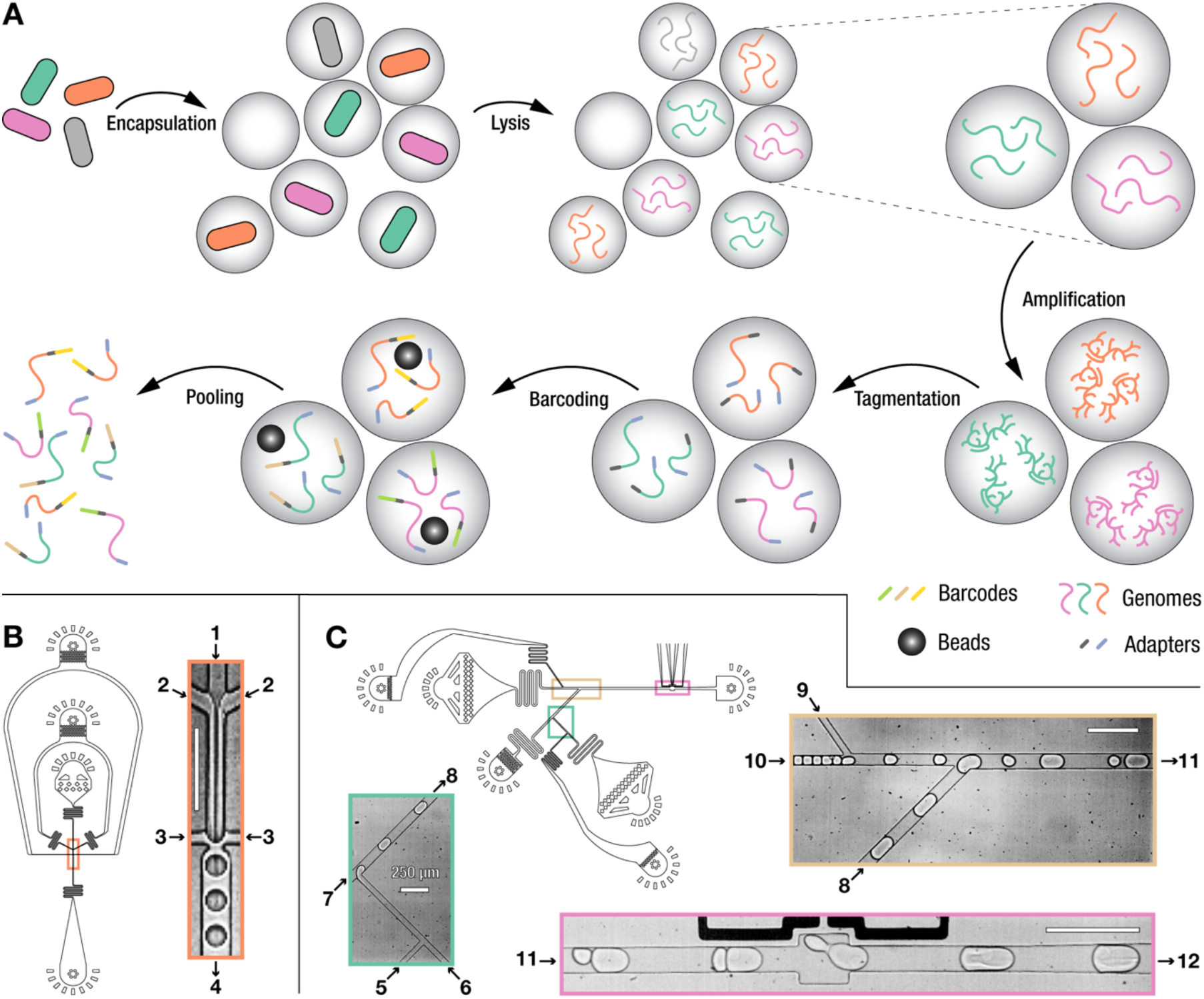
Schematic of the *Microbe-seq* workflow, and microfluidic devices. (A) Schematic of the *Microbe-seq* workflow. Microbes are isolated by encapsulation with lysis reagents into droplets. Each microbe is lysed to liberate the DNA. After lysis, amplification reagents are added to each droplet to amplify the single-microbe genome in each individual droplet. Tagmentation reagents (Nextera) are added into each droplet to fragment amplified DNA and tag them with adapters. Polymerase chain reaction (PCR) reagents and a bead with DNA barcodes are added to each droplet. PCR is performed to label the genomic materials with these primers, and droplets are broken to pool barcoded single-microbe DNA together. (B) Design of the microfluidic device that encapsulates individual microbes into droplets. The orange rectangle outlines the area shown in the image. Labels 1 to 4 are the microbial suspension, lysis reagents, oil, and droplets containing microbes, respectively. Scale bar: 250 microns. (C) Design of the microfluidic device for barcoding that adds beads and PCR reagents to droplets. Colored rectangles outline the areas depicted in the images with corresponding outline colors. Labels 5 to 8 are the PCR reagents, barcoding beads, oil, and droplets each containing a bead and PCR reagents, respectively. Labels 9 to 11 are the oil, sample droplets, and droplet pairs (sample droplet and droplet containing a bead and PCR reagents), respectively. Label 12 are the merged droplets containing the sample, a bead, and PCR reagents. Scale bars: 250 microns.

The raw sequencing data is a collection of DNA sequences; each of these *reads* contains two parts: a barcode sequence shared among all reads from the same droplet, and a sequence from the genome of the microbe originally encapsulated in that droplet. The collection of microbial sequences associated with a single barcode represent a *single amplified genome* (SAG) (Rinke et al., 2014).

### Single-microbe genomics in a community of known bacterial strains

To characterize the nature of the information contained within each SAG, we determine whether each SAG contains genomes from one or multiple microbes, and how much of a microbe’s genome is contained in each SAG. Consequently, we apply our methods to a mock community sample that we construct from strains whose genomes are already known completely, providing an established reference to check the quality of each SAG. The mock sample contains four bacterial strains in similar concentrations, each of which has a complete, publicly-available reference genome: gram-negative *Escherichia coli* and *Klebsiella pneumoniae*, and gram-positive *Bacillus subtilis* and *Staphylococcus aureus*. From the mock sample, we recover 5497 SAGs, each containing an average of 20K reads (Table S1).

To assess the extent to which each SAG contains genomic information from only a single microbe, we compare each of its reads to the reference genomes of the four bacterial strains. We align each read against each genome; i.e., if a genome has a sequence that matches sufficiently closely to a given read, we designate this match as a successful alignment between that read and the genome (Langmead and Salzberg, 2012); we identify the genome that contains the sequence that most closely matches each read as the closest-aligned genome. If a SAG includes reads from multiple microbes, its constituent reads likely connect with a mix of different closest-aligned genomes; by contrast, if the reads from a SAG originate from only one microbe, then those reads will connect to the same closest-aligned genome. To test this, for each SAG, we examine all reads that align successfully to at least one of the four genomes, and determine the percentage of those reads that share the same closest-aligned genome; we define the highest of these four values as the *purity* of that SAG (Lan et al., 2017). Within the mock sample, we find that 84% (4612) of the SAGs have a purity that exceeds 95%, which we designate as high-purity; these data demonstrate that an overwhelming majority of SAGs represent single-microbe genomes, as shown in the distribution in Figure 2A.

**Figure 2.**
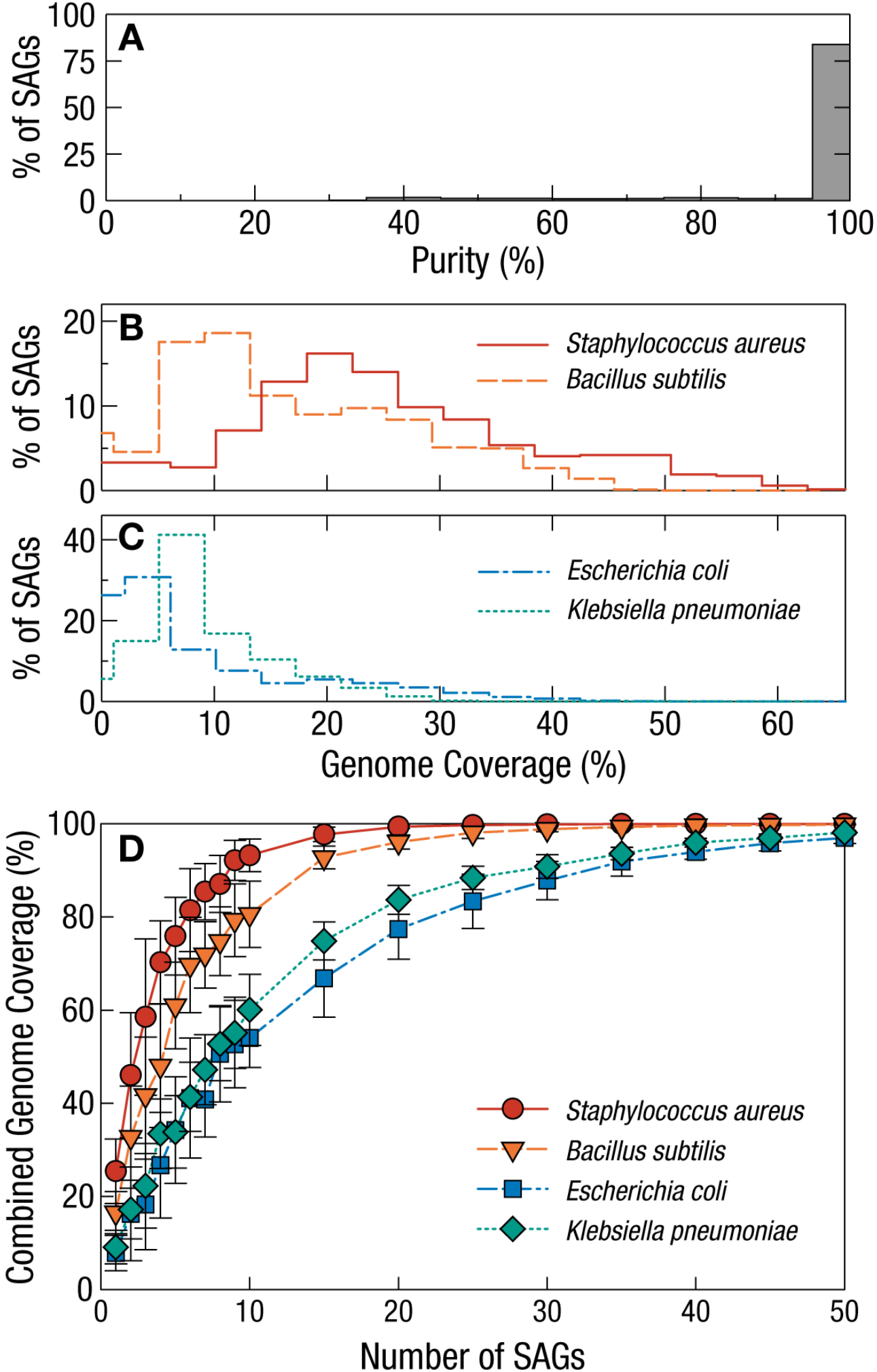
Single-microbe genomics in a mock community sample of known bacterial strains. (A) Purity distribution of all single amplified genomes (SAGs) from the mock community sample, which for the overwhelming majority of SAGs exceeds 95%, demonstrating a single-microbe origin for the DNA in each of these SAGs. (B) Genome coverage rates for SAGs of the two gram-positive bacterial strains, *Bacillus subtilis* and *Staphylococcus aureus*, broadly distributed around average values of 17% and 25%, respectively. (C) Genome coverage rates for SAGs of the two gram-negative bacterial strains, *Escherichia coli* and *Klebsiella pneumoniae*, more narrowly distributed around average values of 8% and 9%, respectively. (D) Combined genome coverage of reads as a function of the number of SAGs where these reads come from; error bars denote standard deviation. In all cases, a few dozen SAGs contain essentially all the information of the microbial genome.

For each of these high-purity SAGs, we identify each base in the corresponding reference genome that has at least one read from that SAG that aligns successfully to it; we use this information to calculate the *genome coverage*, the ratio of these aligned bases to the total number of bases in the reference genome, for each SAG. We find that the genome coverage is broadly distributed around the average values of 17% for *B. subtilis*, and 25% for *S. aureus*, as shown in Figure 2B. The coverage for these gram-positive strains is roughly double that of the coverage for the gram-negative strains, which peaks more narrowly around the average values of 8% for *E. coli* and 9% for *K. pneumoniae*, as shown in Figure 2C (Table S1); the comparatively smaller genome sizes of the gram-positive strains likely contributes to this observed coverage difference.

The genome coverage of each individual SAG is incomplete; one way to overcome this limitation is to combine the genomic information from multiple microbes that belong to the same strain, which are known to share nearly-identical genomes. To explore how the genomic information contained within a group of SAGs depends on the number of SAGs in the group, we select randomly a subpopulation of SAGs from the group that matches each of the four reference genomes, and determine the total combined coverage of all of the reads within that group of SAGs. We calculate the combined coverage as a function of the number of SAGs in that group and find that it increases with SAG group size. While the specific number of SAGs needed to reach any given combined coverage varies between strains, in all cases the information that would be needed to reconstruct essentially-complete genomes is in principle present within any randomly-selected group of several dozen SAGs, as shown in Figure 2D.

### Human gut microbiome samples

To explore the utility of this single-microbe sequencing, we apply the droplet-based approach to a complex microbial community. We explore the human gut microbiome, expected to contain on the order of a hundred species (Human Microbiome Project, 2012a). We examine seven stool samples collected from a healthy individual over a year and a half, for which both shotgun metagenomic data sets and cultured isolate genomes have been reported separately (Poyet et al., 2019). We recover 1000 to 7000 SAGs per sample, for a total of 21,914 SAGs (Table S2). Each SAG contains an average of about 70K reads, so that each sample contains several hundred million reads.

### Microbial diversity in the human gut microbiome

The human gut microbiome comprises a large number of microbial taxa (Lozupone et al., 2012); its diversity is typically assessed with shotgun metagenomics. To assess the diversity measured by the droplet-based protocols, as compared to metagenomic methods, we collect all reads from all SAGs in each sample and ignore the barcode information that connects reads from the same SAG; this yields a data set of similar structure to that used in metagenomics. We classify each read in each sample at the genus level by comparing to the public database of microbial genomes (Wood and Salzberg, 2014); we also perform this genus-level comparison on each read from the corresponding metagenomic data sets. Our stool samples contain a few thousand cells; by contrast, metagenomics typically accumulates the genomic data from millions of cells. Nevertheless, we recover 96.9% to 99.8% of the genera found by metagenomic analysis of the seven stool samples, as shown Figure 3A. We recover similarly high fractions of the taxa identified using metagenomics at broader taxonomic levels (Table S2).

**Figure 3.**
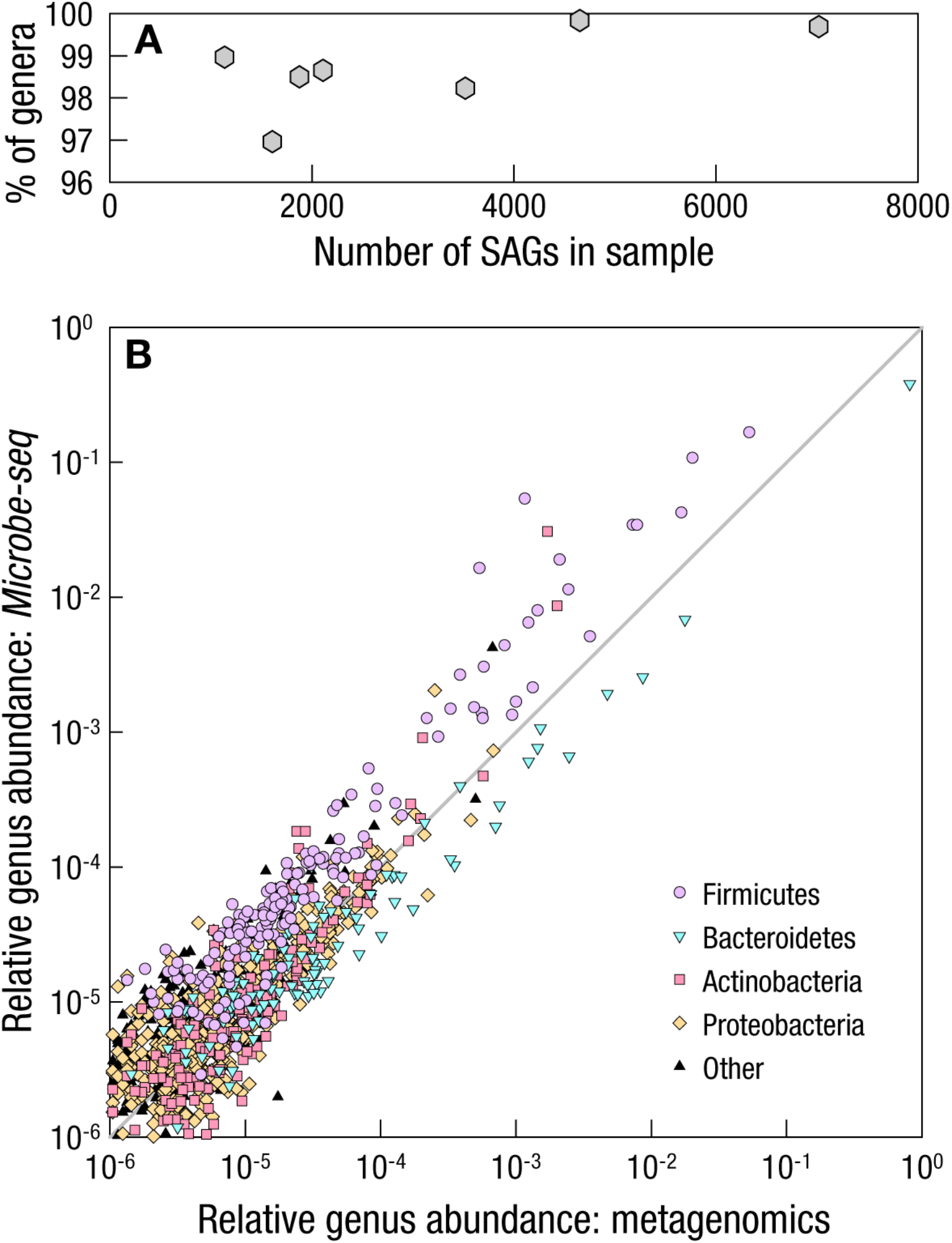
Similar microbial diversity in the human gut microbiome captured both by metagenomics and by *Microbe-seq*. (A) Fraction of genera found by metagenomic analysis that also appear in the corresponding *Microbe-seq* dataset as a function of the number of SAGs in each of the seven samples, which averages 99% and, in all cases, exceeds 97%. These data demonstrate that *Microbe-seq* identifies the presence of a similar range of taxa as metagenomics. (B) Comparison of the relative abundance of genera captured by metagenomics, and by *Microbe-seq*, for the earliest of the seven samples collected. The proximity of the data points to a line of slope 1 on the log-log plot indicates that both methods capture similar abundance.

In addition, we determine for each genus its relative abundance using the droplet-based method and using metagenomics; to compare these results, we plot both abundance values on a scatterplot with equal axes. We find that both techniques report similar values, as indicated by the proximity of the data points to a line of slope 1, as shown for the first stool sample in Figure 3B; we find the same behavior for all other samples (Figure S2). These genera belong to several phyla, predominantly Firmicutes and Bacteroidetes, with additional matches to Actinobacteria, Proteobacteria, and Fusobacteria, consistent with the known composition of the human gut microbiome in industrialized populations (Lloyd-Price et al., 2017); we indicate these phyla with differently-colored symbols in the figure. In addition, we also observe the presence of viruses in the data obtained using the droplet-based approach, consistent with the same observation in metagenomics (Table S2). These data demonstrate that the droplet-based methods capture the same diversity as that captured using metagenomics.

### Genomes of microbial species in the human gut microbiome

To exploit more fully the data from the droplet-based methods, the contents of each SAG must be identified, which is best done by comparison with known genomes. In the case of the mock sample, we identify each SAG by comparing its reads to pre-existing reference genomes. By contrast, in the case of the human gut microbiome samples, no complete set of genomes from all major strains exists, and certain species may not even appear in public reference databases; more generally, it is not possible to identify SAGs from complex microbial communities using comparison with pre-existing reference genomes. Based on the data from the mock sample, we expect the coverage of the SAGs to be far from complete, thereby precluding an individual SAG from being used as a reference genome. Consequently, we develop an approach that does not consult external genomes, but instead combines the genomic information from multiple SAGs to co-assemble genomes, to enable identification of individual SAGs.

The first task in this approach is to identify SAGs that correspond to the same species. Within each SAG, we *de novo* assemble the reads with overlapping regions into *contigs* (Bankevich et al., 2012), longer contiguous sequences of bases; the resulting set of these contigs forms that SAG’s partial *genome*, which we expect from the mock sample to cover only a few percent of the total genome, somewhat less than the coverage of the reads themselves. The overlap between two genomes from the species is expected to be roughly the square of this coverage, generally less than one percent; consequently, any two genomes from SAGs of the same species will likely share only a few or even no direct overlaps. This low overlap prevents direct sequence alignment from being a robust method to determine the similarity of two partial genomes; instead, for each SAG’s genome, we use a hash function to extract a signature that is indicative of the complete genome (Ondov et al., 2016). We compare the signatures of all pairs of genomes, using hierarchical clustering to group SAGs with similar partial genomes into preliminary data bins. For all SAGs within each of these bins, we treat all of the reads equally and *co-assemble* them into that bin’s tentative genome. We then calculate new signatures for the tentative genomes, and re-compare their similarity, iterating this process to consolidate bins that should contain sequences from the same species.

This initial grouping process may generate bins containing reads from multiple taxa. In response, we examine how the reads within each bin align to the contigs in its tentative co-assembled genome. For each contig, we examine the reads that align to that contig successfully; if two different contigs have non-overlapping subgroups of SAGs whose reads align successfully, then each of these subgroups likely correspond to different taxa (Yu et al., 2017). In these cases, we create new bins from these subgroups, and co-assemble their tentative genomes; these genomes should in principle represent only a single taxon.

After this bin-splitting process, multiple bins may contain genomes that correspond to the same species, which we may identify by comparing their genomes. However, in contrast to the earlier steps, each bin at this stage contains a genome co-assembled from many SAGs, which is large enough to share overlapping sequences with genomes from other bins that represent the same species; consequently, we can compare the sequences of tentative genomes directly, without needing to rely on comparatively less-precise hashes. For all pairs of these tentative co-assembled genomes, we calculate their *average nucleotide identity* (ANI), a metric that estimates the similarity of two genomes by comparing their homologous sequences (Jain et al., 2018); we use an ANI value exceeding 95% to indicate that both genomes belong to the same species (Jain et al., 2018). Using this criterion, we merge all bins corresponding to the same species and co-assemble their constituent reads to yield refined genomes of individual species (STAR methods).

To evaluate the quality of each of these refined co-assembled genomes, we count single-copy marker genes to estimate two metrics: *completeness*, the fraction of a taxon’s genome that we recover; and *contamination*, the fraction of the genome from other taxa (Parks et al., 2015). We find that 52 co-assembled genomes have completeness greater than 0.9 and contamination less than 0.05, designated as *high-quality* (Almeida et al., 2019; Almeida et al., 2020; Pasolli et al., 2019); we also find 24 other co-assembled genomes have completeness > 0.5 and contamination < 0.1, designated as *medium-quality*. More than three-quarters (16723) of the SAGs belong to one of these 76 species, demonstrating that we successfully reconstruct reference genomes for the overwhelming majority of SAGs. Furthermore, these 76 genomes include six species with fewer than two dozen SAGs, representing 0.1% of total SAGs, providing strong evidence that we can detect even rare taxa and reconstruct their genomes successfully.

To determine whether each genome corresponds to a single species known to occur in the human gut microbiome, we compare each co-assembled genome against a public database (GTDB-Tk) (Chaumeil et al., 2019), using the ANI > 95% criterion to identify matches of the same species. We find a broad mix of species from diverse phyla (Table S3), including Firmicutes, Bacteroidetes, Actinobacteria, Proteobacteria, and Fusobacteria; in particular, abundant species include several well-known in the human-gut microbiome, including *Faecalibacterium prausnitzii, Bacteroides uniformis*, and *Bacteroides vulgatus*. For each of these 76 genomes, we list the name, colored by corresponding phylum; we illustrate its phylogenetic relationships with other species using a dendrogram; and we indicate the number of SAGs used in its co-assembly with the length of the outer bars, shaded for those of high quality, in Figure 4.

**Figure 4.**
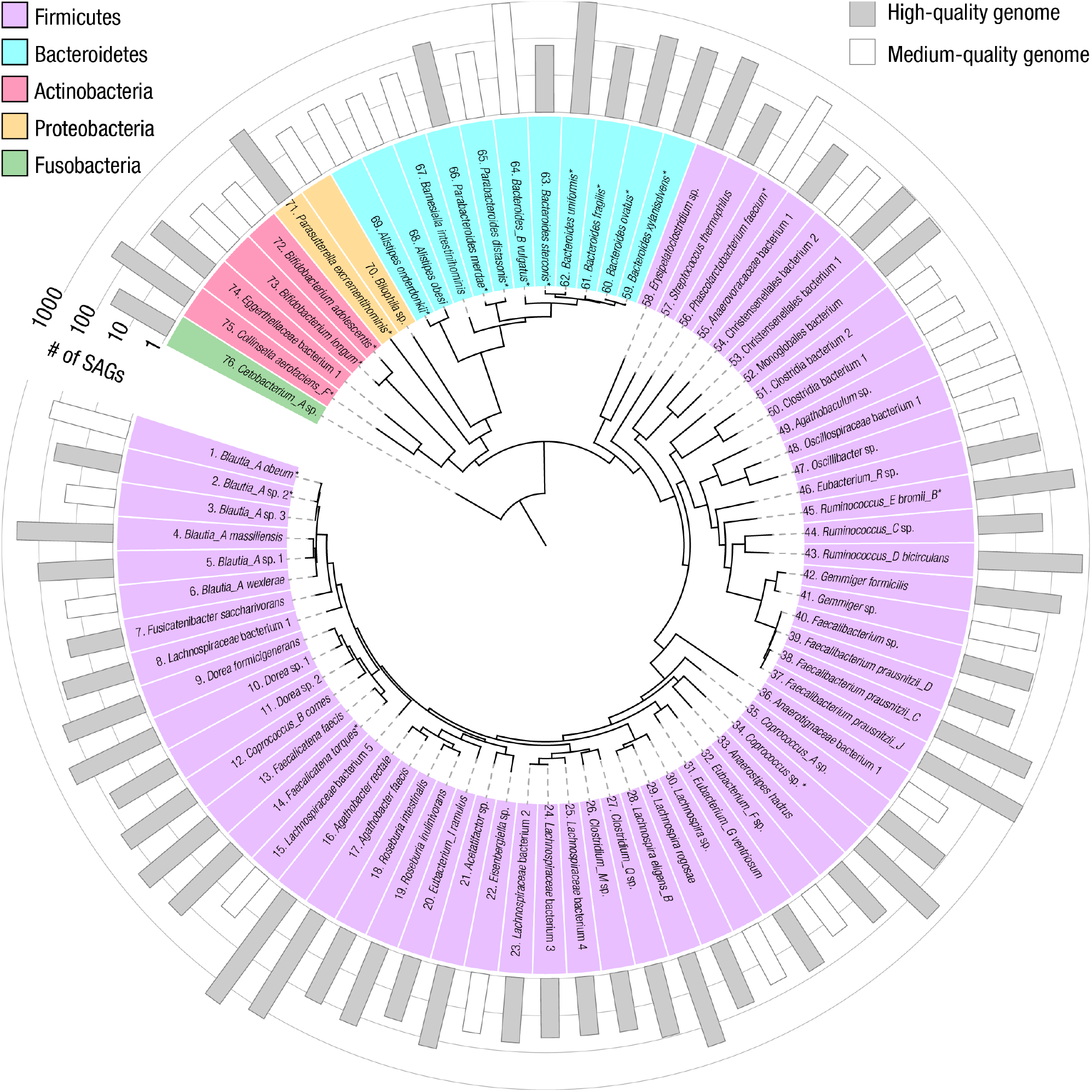
Co-assembled genomes of 76 bacterial species in the human gut microbiome of a single individual. The 76 bacterial species with high- or medium-quality co-assembled genomes. Phylogeny constructed from ribosomal protein sequences represented by the dendrogram in the center of the circle. The phylum of each species is indicated by background color (as in Figure 3) behind each listed species name (GTDB-Tk database), marked with an asterisk for the 19 species with genomes from isolates cultured from the same individual. The number of SAGs used for co-assembly (abundance) is indicated by the bars in the outermost ring, shaded in gray for the 52 high-quality genomes, and unshaded for the 24 medium-quality genomes.

Because there exists for these samples a large number of isolates cultured from the same individual (Poyet et al., 2019), we compare the co-assembled genomes with the “gold-standard” genomes derived from isolates. We find 19 species for which the co-assembled genomes have corresponding isolate genomes, which we mark with an asterisk following each species name in Figure 4. In 18 species, the ANI exceeds 99.5%; these data provide strong evidence for the faithful reconstruction of genomes that closely match those of the cultured isolates, with low contamination.

With only a small set of culture-free experiments, we recover a broad set of accurate reference genomes from more species than those recovered from any other single gut microbiome. These genomes enable us to assign the overwhelming majority of single-microbe SAGs in the sample to one of these 76 species.

### Strain-resolved genomes in the human gut microbiome

Many species in the human gut microbiome are represented by multiple strains (Garud et al., 2019); different strains may play distinct roles within complex microbial communities and express different sets of genes to carry out these roles (Albanese and Donati, 2017). Linking specific genes, and consequently their functionality, to the strains which contain them requires knowledge of the genomes from those individual strains. Moreover, because each microbe inherently represents only a single strain, definitive identification of each SAG requires strain-resolved reference genomes.

To explore the possibility that our co-assembled genomes contain contributions from more than a single strain, we examine further the comparison between the 19 co-assembled genomes and cultured isolates of the same species; each of these isolates represents only a single strain. In general, the co-assembled genome of a species with multiple strains contain some contigs unique to each strain; not all of these contigs appear in the single-strain genomes of the corresponding isolates. Consequently, we determine the *shared genome fraction*, the percentage of bases in each co-assembled genome that are shared with isolate genomes from the same species. We find that for the comparison in 16 species, the shared genome fraction is above 96%, and the ANI value exceeds 99.9%; these data suggest that each of these 16 co-assembled genomes represents a single strain. By contrast, for the remaining three species, *Blautia obeum, Bacteroides vulgatus*, and *Parasutterella excrementihominis*, the shared genome fraction is far lower, between 70% and 90%, and ANI are all less than 99.6% (Figure S3); these low values suggest that the genomes of these three species include multiple strains or strains that do not appear among the cultured isolates. In principle, comparing directly all pairs of SAGs to estimate the fraction of their shared genomes could distinguish strains. However, the coverage of each SAG is expected to be on average less than 25%, for example 7% of the genome for the *Bacteroides vulgatus*; this coverage suggests that such pairwise comparisons will not in general be reliable and instead motivates a different approach.

To distinguish strains, we develop a method that leverages the differences among homologous sequences between SAGs, specifically, the single-nucleotide polymorphisms (SNPs). To illustrate this method, we examine ∼900 SAGs of *Bacteroides vulgatus*, the most abundant of the three species; we align reads from each SAG against the co-assembled *Bacteroides vulgatus* genome and identify ∼12000 total SNP locations. For each SAG, we determine the *SNP coverage*, the fraction of all SNP locations in the genome that occur among the reads of that SAG; this SNP coverage is, on average, 8%, comparable to the average genome coverage. For each pair of SAGs, we measure the fraction of total SNP locations that occur in both, and find this fraction to be about 0.7%, corresponding to about 80 SNPs, consistent with roughly the square of the SNP coverage. Microbes of the same strain have nearly-identical genomes (Poyet et al., 2019; Zhao et al., 2019), so that two SAGs representing the same strain almost always have the same base at each SNP location shared by both SAGs; conversely, SAGs representing different strains show significantly lower similarity (Garud et al., 2019). Inferring the similarity of the bases at shared SNP locations in each pair of SAGs is governed by a binomial process; therefore, the average of 80 SNPs in each SAG pair should be sufficient for a robust inference, with uncertainty of a few percent or less. Consequently, the comparison of SNPs provides a promising approach to determine strains.

To test this possibility, we examine in all pairs of SAGs the bases at all shared SNP locations and determine the fraction of these locations where both SAGs have the same base. To probe whether these SAGs fall into any distinct groups, we visualize the SNP similarity between all pairs of SAGs with dimensional reduction (Becht et al., 2018). Strikingly, we find that the SAGs fall into four clearly distinct clusters, as shown in Figure 5A. We validate independently the presence of these SAG groups with hierarchical clustering, which yields the same groupings with 99.8% overlap.

**Figure 5.**
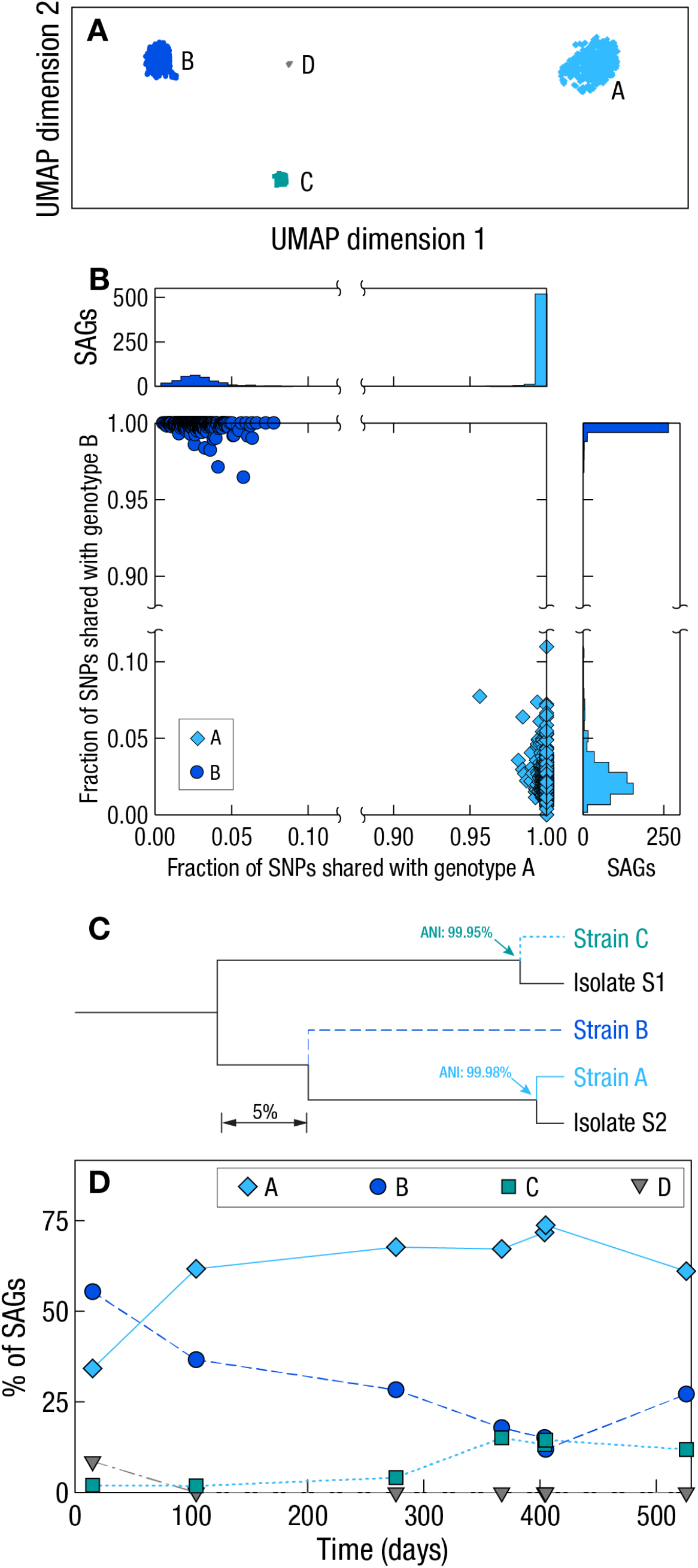
Strain-resolved genomes of *Bacteroides vulgatus* in the human gut microbiome. (A) Dimension-reduction (UMAP) visualization of *Bacteroides vulgatus* SAGs, based on comparison of their sequences at single nucleotide polymorphism (SNP) locations. SAGs fall into four distinct, widely-separated clusters; the symbol for each SAG is colored according to the cluster in which it is grouped. (B) Scatterplot and histograms illustrating the fraction of SNPs from each SAG that match consensus genotypes A and B, for SAGs in these two most-abundant clusters. In almost all cases, each SAG shares the same base in more than 99% of the SNP locations in its corresponding consensus genotype; by contrast, the SNP overlap with the consensus genotype of the other cluster is much lower, typically only a few percent. Symbols in each cluster colored as in (A). (C) Phylogeny of the co-assembled high- and medium-quality genomes of *Bacteroides vulgatus* strains, and comparison with the corresponding genomes of strains of isolates cultured from the same individual. The horizontal axis of the dendrogram represents the fraction of shared sequences. Average nucleotide identity (ANI) values between these strain-resolved genomes are labeled in the plot, demonstrating that co-assembled strain C and isolate S1 are the same strain; similarly, co-assembled strain A and isolate S2 are the same strain. By contrast, the second most-abundant strain, B, does not appear among the isolates cultured from the same individual. (D) Relative abundance of the four *Bacteroides vulgatus* strains in the seven longitudinal samples, which appears to change gradually over time.

To test whether these clusters correlate with different strains, we examine within each SAG cluster the bases at SNP locations. We determine which base occurs most frequently at each SNP location; the set of these bases at each SNP location forms the *consensus genotype* of each SAG cluster. Then, for each SAG, we calculate the fraction of its SNPs that have the same base at the corresponding location in the consensus genotype of each of the four SAG clusters. Within each SAG cluster, we find that constituent SAGs share extremely high SNP similarity with the corresponding consensus genotype: for example, in the two clusters with the highest number of SAGs, almost all SAGs have the same base in more than 99% of the SNP locations, as shown in the scatterplot and histograms in Figure 5B. By contrast, SAG clusters show much lower overlap with the consensus genotypes of other clusters; for the two clusters with the highest number of SAGs, all SAGs in each cluster share fewer than 10% of the bases at SNP locations with the consensus genotype of the other cluster, as shown in the figure. These trends persist among the other clusters (Figure S4). Together, these results provide strong evidence that SAGs within these clusters represent the same strain.

To examine further whether these four clusters correspond to actual *Bacteroides vulgatus* strains, we co-assemble the reads within each SAG cluster. We obtain high-quality genomes for the two groups with the most SAGs, which we label candidate strains A and B; one medium-quality genome, C; and one additional genome of lower quality, D (Table S4). We compare these co-assembled genomes with the genomes of two distinct *Bacteroides vulgatus* isolate strains cultured from the same individual (Poyet et al., 2019). Strikingly, we find that both isolate genomes have closely-matching co-assembled counterparts, A and C, with ANI values and shared genome fractions exceeding 99.9% and 97%, respectively, as shown in Figure 5C. These high values are consistent with those that occur between genomes of the same strain, thereby providing strong evidence that these co-assembled genomes each represent a single, genuine strain of *Bacteroides vulgatus*. Surprisingly, the second-most populous cluster, candidate strain B, with several hundred SAGs, does not appear among the nearly one hundred isolates of *Bacteroides vulgatus* cultured from the individual (Poyet et al., 2019). Together, these results demonstrate the capabilities of this SNP-based approach to identify correctly both the major known strains of *Bacteroides vulgatus* and potentially new strains that have not been cultured, while at the same time enabling the accurate co-assembly of their genomes.

We further apply this SNP-based analysis to the remaining 29 species with high-or medium-quality species-level genomes, each co-assembled from more than 100 SAGs. We find five additional species with multiple strains, co-assemble their genomes (Figure S5 and Table S4), and compare to corresponding isolate genomes cultured from the same individual. We find excellent agreement for *Blautia obeum*, with an ANI of 99.9% and shared genome fraction of 95%, again confirming, just as in the case for *Bacteroides vulgatus*, that our co-assembled genome represents a single, genuine strain. In total, we obtain 84 high- and medium-quality strain-resolved genomes from 76 species, from just one set of experiments. Remarkably, we are able to achieve this accurate identification of strains and the co-assembly of their genomes, even with a level of coverage that yields an average of less than a hundred shared SNP locations between all pairs of SAGs.

The capability to identify the strain of each individual SAG also enables us to follow the relative abundances of these strains over time in this individual, providing insight on the population dynamics. The abundances of these strains appear to shift only gradually throughout the year and a half over which samples were collected; for instance, we observe quite similar abundances in *Bacteroides vulgatus* in the two samples collected on successive days around day 400, as shown in Figure 5D.

The results demonstrate the capability of this approach to resolve sub-species strains and reconstruct their strain-resolved genomes, from species with more than 100 SAGs, even when the SAGs have coverage only of around 10% of the genome. Furthermore, this droplet-based approach can obtain strain-resolved genomes from strains which have not been cultured; this is of particular significance to the human gut microbiome, where many strains are difficult to culture. Consequently, this method contributes a new way to examine the strain-resolved structure and dynamics of the genomic information within the human gut microbiome that is independent of the bias imposed by what has been cultured. These high-quality, strained-resolved genomes from a broad range of strains from the gut microbiome of a single person not only allow greater precision in the identification of the overwhelming majority of SAGs, but further enable the probing of broader genomic aspects of the microbial community, particularly those involving microbes of different strains.

### Horizontal gene transfer (HGT) within the human gut microbiome

One particularly exciting genomic aspect of microbial communities is how microbes exchange genetic information; one of the most well-known mechanisms is horizontal gene transfer (HGT), which has been observed to be frequent within the human gut microbiome (Hehemann et al., 2010; Huddleston, 2014; Keeling and Palmer, 2008; Smillie et al., 2011). The large number of strain-resolved genomes originating from the gut microbiome of a single human offers the intriguing potential to detect comprehensively and accurately the sequences shared between specific microbial taxa, a hallmark of HGT.

One method to establish HGT between two different microbial species identifies common sequences that occur in the genomes of both, and often utilizes cultured isolates (Groussin et al., 2020; Smillie et al., 2011). Reads from each cultured isolate descend ultimately from a single microbe; consequently, these reads are usually connected to the genome of a single strain. By contrast, metagenomic reads originate from multiple different species and strains, to which the short reads cannot usually be connected, thereby making the determination of HGT with metagenomics alone extremely challenging. In the present work, the ability to link each read to a single SAG offers a possible way to overcome this challenge, by identifying sequences common to strain-resolved genomes of different species, without the uncertainty as to the strains in which they occur.

To assess the effectiveness of this approach of detecting HGT in a single person, we identify common sequences between all pairs of the 13 high-quality strain-resolved co-assembled genomes, for which we also have corresponding isolate genomes, from the same individual. We identify 19 HGT pairs among these 13 strain-resolved co-assembled genomes; applying the same analysis to the 13 corresponding cultured isolate genomes, we also identify those same pairs, plus one additional pair, as shown in the correlation matrix in Figure 6A. These data correspond to a 95% rate of agreement, with no false positives, demonstrating the sensitivity and specificity of this approach.

**Figure 6.**
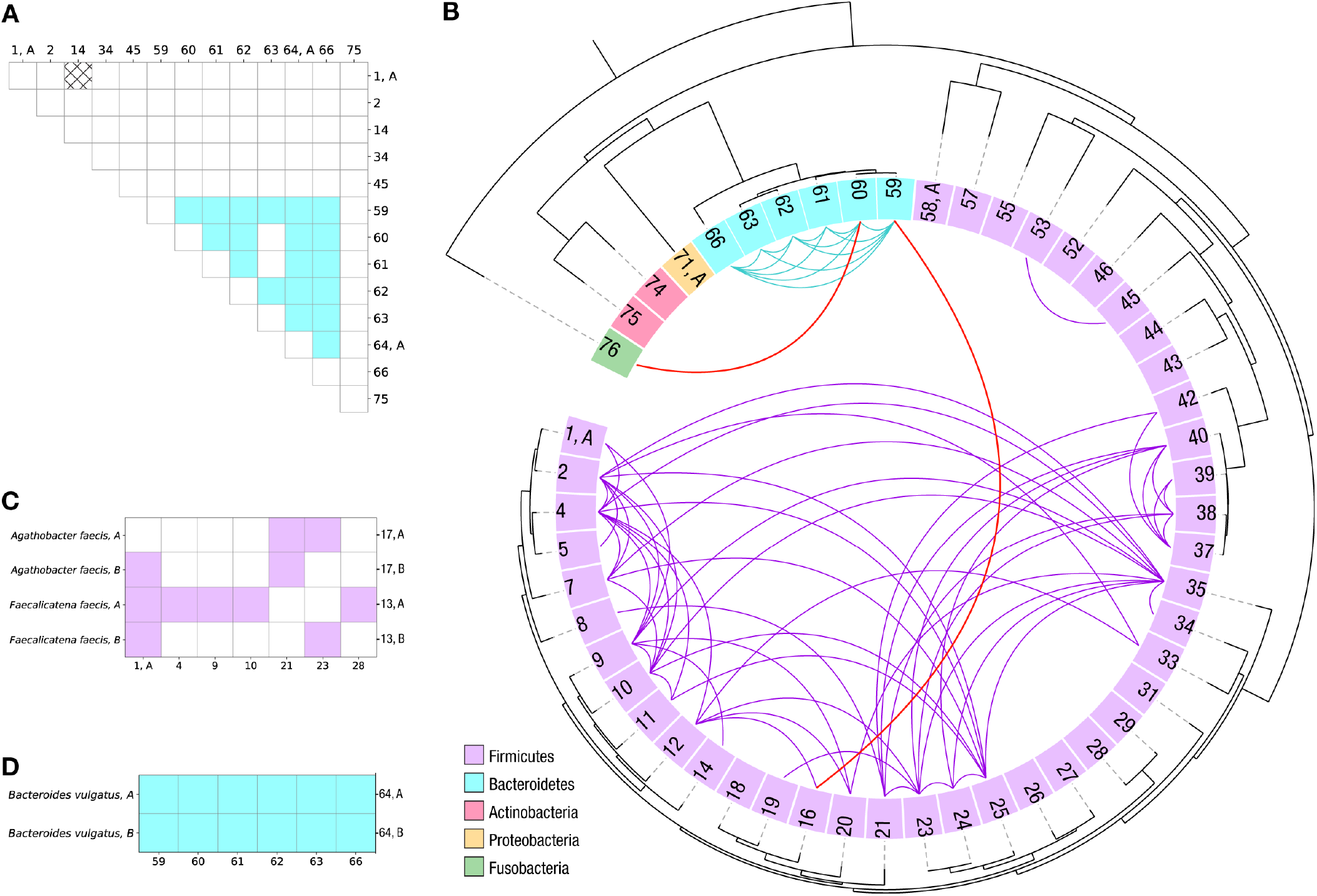
Horizontal gene transfer (HGT) among bacterial strains within the human gut microbiome of a single individual. (A) Comparison of HGT within 13 high-quality single-strain co-assembled genomes, and, separately, within the 13 corresponding genomes of isolates cultured from the same individual. Numbers along both axes are the labels indicating the species, as in Figure 4. Cells filled with solid blue represent the 19 pairs of genomes in which HGT is detected between them, both within the co-assembled data sets and within the isolate data sets. The cell with the black crosshatch pattern represents the single genome pair where HGT is detected only between the cultured isolate genomes and is not observed between the corresponding pair of co-assembled genomes. Unfilled cells represent pairs of genomes in which no HGT is detected in either dataset. (B) HGT among the 51 species with a single high-quality strain-resolved genome, following the order, numbers and colors of Figure 4. Detected HGT between two genomes is indicated with a curve whose color matches that of the phylum of each pair, in cases where both are within the same phylum; red curves denote the cases in which the genomes are in different phyla. The overwhelming majority of detected HGT occurs between two species within the same phylum. (C)-(D) HGT between the species in B, and different strains of the same species, for bacterial species with multiple high-quality strain-resolved genomes. (C) For the bacteria in phylum firmicutes, *Agathobacter faecis* and *Faecalicatena faecis*, each strain has HGT with different sets of species. (D) For the phylum Bacteroidetes, the only multi-strain species is *Bacteroides vulgatus*, which has HGT between both of its strains and all other species in this phylum.

With this quantitative validation, we investigate HGT among the 51 species, each with a high-quality genome that represents a single strain; these include 41 species in the phylum Firmicutes, 6 in Bacteroidetes, 2 in Actinobacteria, and 1 each in Proteobacteria and Fusobacteria. This yields 836 pairs of species where both are within the same phylum: 820 within Firmicutes, 15 within Bacteroidetes, and one within Actinobacteria; among these, we detect HGT between 68 pairs of species, as shown in Figure 6B. By contrast, among the 439 pairs of genomes from different phyla, we detect HGT between only two species pairs: one between Bacteroidetes and Firmicutes, and another between Bacteroidetes and Fusobacteria, as shown in Figure 6B. We also detect HGT involving those species with multiple high-quality strain-resolved genomes: *Agathobacter faecis* and *Faecalicatena faecis* and *Bacteroides vulgatus*. Interestingly, we observe that individual strains of *Agathobacter faecis* and *Faecalicatena faecis* exchange genes with different Firmicutes species, as shown in Figure 6C; by contrast, both strains *Bacteroides vulgatus* undergo HGT with the same set of other Bacteroidetes species, as shown in Figure 6D.

These data are consistent with observations that HGT in the gut microbiome occurs largely within phylum-level gene pools (Jiang et al., 2019). Furthermore, the differences in HGT between and within Firmicutes and Bacteroidetes suggest that the ways in which bacteria import genetic material may not be universal, but instead might have some phylum-dependent factors. These observations demonstrate the importance of having a large number of strain-resolved, high-quality genomes to understand the nature of HGT within the human gut microbiome.

### Host-phage association in the human gut microbiome

Our approach is not limited to only bacteria, but also applies to other types of microbes. Indeed, the diversity analysis reveals the presence of viruses, specifically crAssphage, the most-abundant bacteriophage at present recognized from the human gut microbiome (Guerin et al., 2018; Siranosian et al., 2020). The general regulatory role of bacteriophages, thought to modulate the abundance and behavior of bacteria, is only beginning to be understood within complex microbial communities (Divya Ganeshan and Hosseinidoust, 2019; Sutton and Hill, 2019). The droplet-based method encapsulates not only an individual bacterium, but also any bacteriophages physically co-located with it, providing a direct means to probe *host-phage association*. To explore this association, we compare the reads in each SAG to the crAssphage genome; we find that a few dozen SAGs contain a significant fraction of crAssphage-aligned reads. Moreover, many of these SAGs also contain a significant fraction of reads which do not align to the crAssphage genome, but instead to bacterial taxa; we align these reads against the co-assembled genomes of 76 species to identify which, if any, bacterial species might associate with crAssphage strain in this particular individual.

Significantly, we find that 14 SAGs are associated with only one species, *Bacteroides vulgatus* (*p*-value = 4*10^−9^, Fisher’s exact test) (Table S5), and that no other species associates significantly with crAssphage, as shown in Figure 7A. These data strongly suggest *Bacteroides vulgatus* as the *in vivo* host species for crAssphage in this individual, consistent with previous evidence that crAssphage is likely to be associated with *Bacteroides* species (Guerin et al., 2018; Shkoporov et al., 2018); the statistical significance of the association indicates that this is not a result of simple random co-encapsulation.

**Figure 7.**
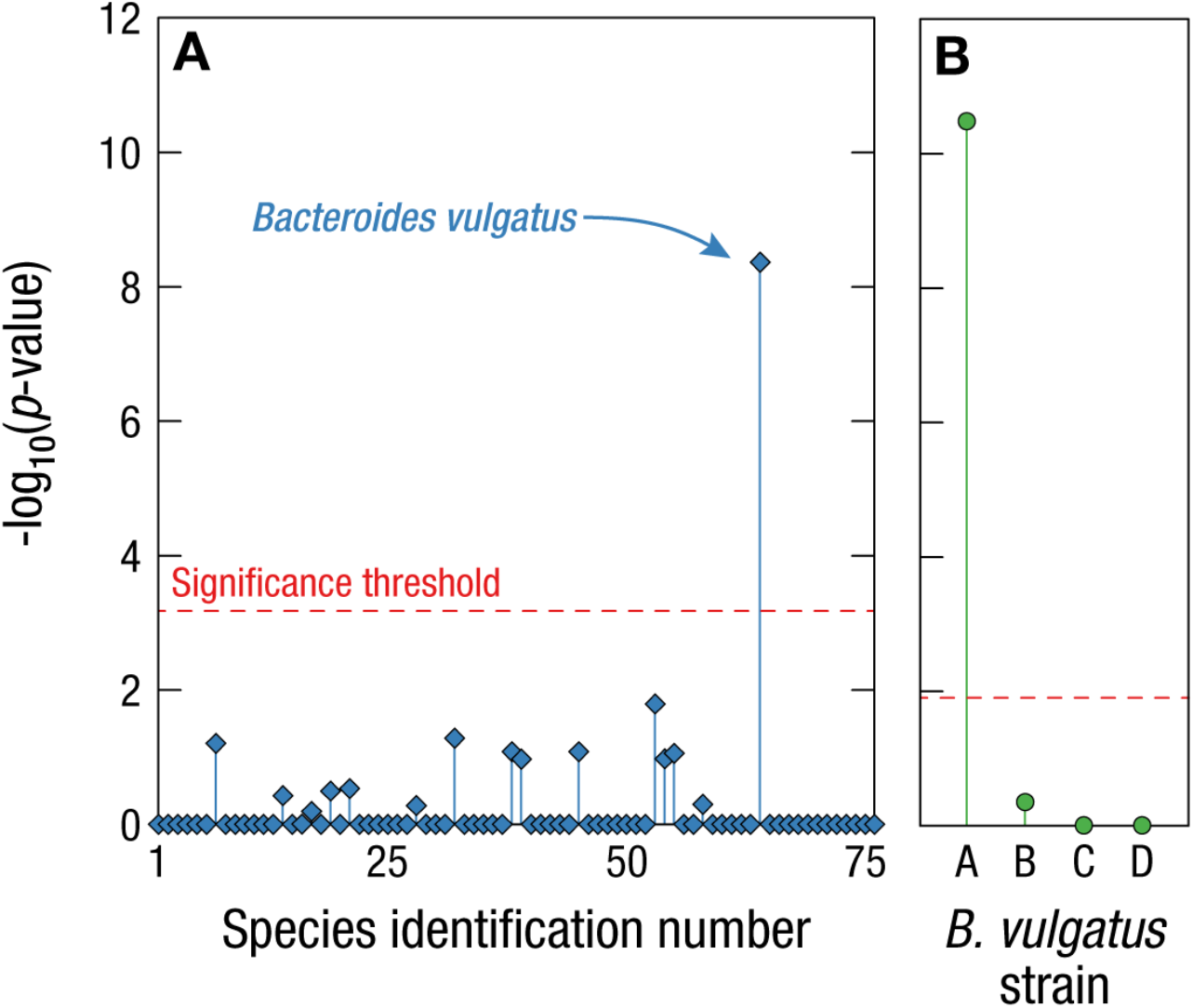
Host-phage association with strain specificity in the human gut microbiome. (A) Association between the bacteriophage crAssphage and bacterial species with high- or medium-quality genomes, with species numbers as in Figure 4. All *p*-values are calculated with one-sided Fisher’s exact test. The only bacterial species that is significantly associated with crAssphage is *Bacteroides vulgatus*. (B) Association between the four strains of *Bacteroides vulgatus* and crAssphage. Only one specific strain of *Bacteroides vulgatus*, the most-abundant strain A, is significantly associated with crAssphage.

Furthermore, the unambiguous assignment of each SAG to one of the multiple strains of *Bacteroides vulgatus* enables even more precise characterization of *in vivo* host-phage association, to the level of specific bacterial strains. Strikingly, we find that 13 SAGs represent the single *Bacteroides vulgatus* strain A, the most abundant (*p*-value=3*10^−11^), as shown in Figure 7B.

These data demonstrate the unique advantages of the droplet-based approach to establish accurately *in vivo* host-phage association not only to an individual species, but even more precisely to a specific strain. We identify which bacterial strains interact with bacteriophages, and which strains do not; the genomic differences between these strains provide preliminary data that may contribute to understanding the molecular mechanisms underlying these host-phage interactions and their longitudinal dynamics in the human gut microbiome.

## Discussion

Using *Microbe-seq*, a high-throughput method combining experiment and computation for single-microbe genomics, we obtain without culturing the genomes of tens of thousands of individual microbes and *de novo* co-assemble the strain-resolved genomes from 76 species, most of which have not been cultured---more strain-resolved genomes than any other single experiment has recovered from a single gut microbiome. Our high-throughput microfluidics-based approach makes practical the individual examination of a sufficient number of microbes to achieve these results, even with an average coverage less than a quarter of the genome. The close agreement with strains where we have corresponding cultured isolates confirms the accuracy of our approach.

These strain-resolved genomes enable the reconstruction of an HGT network within a single human; when sampled over time, these data may allow the monitoring of microbe response, at the level of specific genes in specific strains, to selective pressures unique to that individual, such as disease, diet or antibiotic treatment. In addition, our *in vivo* association between specific strains of bacteriophages and bacteria provide specific starting points to investigate how phages modulate microbial composition and guide subsequent development of phage-based therapeutics.

*Microbe-seq* provides a particularly effective and practical approach, in a single laboratory-scale experiment, to identify and sequence fully all of the major strains in microbial communities beyond the human gut microbiome, without any *a priori* knowledge of constituent microbes. The protocol integrates proven droplet microfluidic operations, applied previously in other contexts, representing a roughly hundred-fold improvement in throughput and cost compared to single-cell sequencing approaches utilizing well plates. Such practical improvements may make feasible the investigation of microbial communities that affect the environments, lives and health of human communities that otherwise lack access to the resources to even begin to investigate these effects.

## Supporting information

microbeseq_methods

microbeseq_supfig

microbeseq_tables

## Acknowledgements

We thank members of the Weitz lab and Alm lab for helpful discussions and Yamei Cai, Wei Chen, Zeyu Chen, Naiwen Cui, Lei Dai, Ruihua Ding, Perry Ellis, Zhifei Ge, Jingjing Gong, Fangqiong Ling, Bingxu Liu, Han Liu, Hao Pei, Raoul Rosenthal, Jizhou Tang, Yongcheng Wang, Jing Xia, Xiaoqian Yu, Zhengrun Zhang, Zeda Zhang, and Zhilun Zhao for general discussions and comments on the manuscript. We thank OpenBiome for providing stool samples. This work was supported at MIT by the Center for Microbiome Informatics and Therapeutics and the Department of Energy (Ecosystems and Networks Integrated with Genes and Molecular assemblies [ENIGMA]), and at Harvard University by the NSF (DMR-1708729) and through the Harvard University Materials Research Science and Engineering Center (DMR-2011754) and the NIH (P01HL120839, R21AI125990, R21AI128623 and R01AI153156) and NASA (NNX13AQ48G, 80NSSC19K0598)).

## Author contributions

W.Z., S.Z., H.Z., P.J.L., E.J.A., and D.A.W. conceived and designed the method; W.Z. developed and performed the experiments with assistance from S.Z.; S.Z., W.Z., Y.Y., D.M.N., and C.L.D. performed data analysis with input from all authors; P.J.L., W.Z., and S.Z. wrote the initial manuscript; D.A.W., P.J.L., W.Z., S.Z., and E.J.A. revised the manuscript; all authors read and commented on the manuscript; E.J.A. and D.A.W. supervised the study.

