## Supplementary material for "*Microbe-seq*: high-throughput, single-microbe genomics with strain resolution, applied to a human gut microbiome": microbeseq_methods

### EXPERIMENTAL MODEL AND SUBJECT DETAILS

We obtain stool samples that are collected by OpenBiome, a non-profit stool bank, under a protocol approved by the institutional review boards at MIT and the Broad Institute (IRB protocol ID # 1402006212). The subject is a healthy male screened by OpenBiome to minimize the potential of carrying pathogens. The person was 28 years old at initial sampling. The subject has been de-identified before receipt of samples. For each experiment, we wash 1-3 uL of stool sample in 1 mL 1X PBS three to five times and resuspend it in 1X PBS with 15% (v/v) Optiprep density gradient medium (Sigma-Aldrich D1556) as the microbial suspension.

### METHOD DETAILS

#### Mock community

We culture four bacteria strains, *Bacillus subtilis* ATCC 6051-U, *Escherichia coli* ATCC 25922, *Klebsiella pneumoniae* ATCC 35657, and *Staphylococcus aureus* ATCC 6538 in 1 mL LB liquid medium (L3522 Sigma Aldrich) overnight. We wash each bacterial culture with 1 mL 1X PBS three to five times and resuspend bacteria in 1X PBS with 15% (v/v) Optiprep density gradient medium (Sigma-Aldrich D1556). We combine approximately the same volume of these four bacterial strains and dilute to a final concentration of 5-50 million microbes/mL.

#### Microfluidic device fabrication

We make our device designs (Figure S1), print them as photomasks (CAD/Art Services, Inc.), and fabricate devices following procedures established in previous reports (McDonald et al., 2000). In brief, we use photolithography and the photomasks to transfer each device design to a silicon wafer with SU8 photoresist. We cast polydimethylsiloxane (PDMS) (Sylgard 184) on the SU8 structure, where the SU8 structure on silicon wafer serves as a master for replica molding. We bake at 65 ºC for at least 2 hours to cure the PDMS and delaminate the resulting PDMS replicas off the master. We seal with glass slides (Corning, 2947) to make microfluidic devices and make their surfaces hydrophobic by flowing Aquapel (PGW Auto Glass, LLC) through the channels. To remove excess residual Aquapel, we flow compressed air in the channels of microfluidic devices and bake the devices at 65 ºC overnight.

#### Isolation and lysis

We isolate microbes by encapsulating them into droplets with lysis reagents using a microfluidic device (Figure S1A and video S1). We put the microbial suspension in a 1 mL syringe (BD Luer-Lok™ 1-mL syringe, 309628) and connect the syringe to microbial suspension inlet through a needle (BD Precisionglide® syringe needles, Z192384-100EA, Sigma Aldrich) and polyethylene tubing (BB31695-PE/2, Scientific Commodities, Inc.). Similarly, we connect lysis reagents and oil, 2% (w/v) surfactant (RAN biotechnologies, 008-FluoroSurfactant) in HFE 7500 (3M), to the device. We remove most of the oil from the bottom of the tube and incubate to lyse the microbes inside droplets.

The incubation program for lysis is: 37 ºC for 30 minutes, 75 ºC for 15 minutes, 95 ºC for 5 minutes and sample storage at 4 ºC.

**Whole genome amplification**

We transfer the droplet emulsion to a syringe and reinject droplets into a microfluidic merger device (Figure S1B and video S1). In the same device, we use a separate droplet maker to form droplets that encapsulate multiple displacement amplification (MDA) reagents. We synchronize the frequency of sample droplet re-injection and reagent droplet-making to form droplet pairs. Applying electric fields of 50-200 V at a frequency of 25 KHz through a pair of electrodes, we merge each droplet pair to add MDA reagents. We incubate to amplify microbial genomes.

The incubation program for MDA is: 30 ºC for 6-8 hours, 65 ºC for 10 minutes and sample storage at 4 ºC.

**Tagmentation**

We merge sample droplets with droplets containing commercially-available tagmentation reagents (Nextera), utilizing a different droplet merger device (Figure S1C and video S1).

The incubation program for tagmentation is: 55 ºC for 10 minutes, and sample storage at 10 ºC.

**Bead synthesis**

We synthesize beads used for barcoding by adopting a previously-reported method (Klein et al., 2015; Zilionis et al., 2017). In brief, we make droplets containing acrydite-modified DNA oligos using a photo-cleavable linker (IDT) and acrylamide:bisacrylamide solution. We keep these droplets at 65 ºC overnight to polymerize them into uniform soft gel beads covalently bonded to the DNA oligos by photo-cleavable linkers. We extend DNA oligos on beads enzymatically with a two-step split-and-pool synthesis protocol to prepare beads with a diverse barcode sequence library. At the first split-and-pool synthesis step, we evenly split beads into a 96-well plate where each well contains a unique barcode-1 oligo (IDT, Table S6). We anneal these oligos with hydrogel oligos and extend them with Bst 2.0 DNA polymerase (M0537L, NEB). After the first split-and-pool synthesis step, we pool beads, wash them and evenly split them into a 384-well plate where each well contains a unique barcode-2 oligo (IDT, Table S6). We perform the second barcode strand synthesis in the same way as we extend the first barcode strand. We avoid exposing beads to strong light.

Each soft gel bead has millions of primers with the same sequence. Each full sequence contains two barcode regions: the first region has a diversity of 96; the second region, 384. Overall, the barcoding bead library has 36864 (96*384) possible sequences. In theory, we can increase the diversity of the library by adding more barcode regions to the barcode sequence or by using more sequences in the two barcode regions.

**Bead preparation for barcoding**

We wash 200 uL of beads with 1 mL bead wash buffer (10 mM pH 8.0 Tris-HCl, 0.1 mM EDTA and 0.1% (v/v) Tween-20), three times in a tube. We withdraw supernatant from the top, leaving 500 uL in the tube. We add 300 uL water and 200 uL 5X Phusion HF detergent-free buffer (F520L, Thermo Fisher) to the tube. We vortex the beads and keep them at room temperature for 1 min. We centrifuge beads, remove supernatants, and use these beads for barcoding.

**Barcoding**

We merge sample droplets with droplets containing PCR reagents and a barcoding bead, using a droplet merger device (Figure S1D and video S1).

The incubation program for barcoding is: 72 ºC for 4 minutes, 98 ºC for 30 seconds; 10 cycles of 98 ºC for 7 seconds, 60 ºC for 30 seconds and 72 ºC for 40 seconds; 72 ºC for 5 minutes, and sample storage at 4 ºC. We use slow ramping of 2 ºC/s at this step.

We observe the merger of some droplets after PCR, possibly during the high-temperature stage of PCR. These droplets are larger and may also contain DNA from multiple microbes. We use a droplet size filter device (Figure S1E) to remove most of these droplets, using a flow rate of 120 uL/h for sample droplets and 2 mL/h for 2% (w/v) oil.

**Droplets pooling and sequencing library preparation**

We first break the emulsion of droplets by adding 200 uL 20% (v/v) PFO (1H,1H,2H,2H-Perfluoro-1-octanol, 370533 Sigma Aldrich) in HFE 7500 (3M) into each sample after PCR. We purify the aqueous phase with 1.1X volume AMPure beads (A63881, Beckman Coulter) and resuspend it into 32 uL DNA suspension buffer (10 mM pH 8.0 Tris-HCl and 0.1 mM EDTA). We use PCR to add sequencing adapters for sequencing (Illumina) and a sample index (Nextera index) to each purified DNA sample so we can sequence multiple samples in one sequencing run.

We prepare a 50 uL PCR mix for each experiment: 2.5 uL water, 10 uL 5X Phusion HF detergent-free Buffer (F520L, Thermo Fisher), 1 µl 10 mM dNTPs (diluted from 25 mM dNTP mix, Thermo Fisher, R1121), 2 uL 10 uM P5PE1 primer (ordered from IDT), 2 uL Nextera i7 primer (Illumina), 0.5 µl Phusion high-fidelity DNA polymerase (F530L, Thermo Fisher), and 32 uL DNA sample in DNA suspension buffer.

The incubation program for PCR is: 98 ºC for 30 seconds; 5-10 cycles of 98 ºC for 7 seconds, 60 ºC for 30 seconds, and 72 ºC for 40 seconds; in the end, 72 ºC for 5 minutes and sample storage at 4 ºC.

We purify samples with 0.8X volume AMPure beads (A63881, Beckman Coulter) and resuspend DNA products into 20 uL DNA suspension buffer (10 mM pH 8.0 Tris-HCl and 0.1 mM EDTA). We store these products at -20 ºC before sequencing.

**Illumina sequencing:**

We sequence at a depth that is about 10-200 thousand reads for each microbe. A custom read-1 primer (IDT) is required for the sample to be sequenced. For a 100 base pair (bp) sequencing run, we use the following sequencing length configurations: read-1 sequence: 45 bp, which only contains barcode sequence; index-1 sequence: 8 bp; read-2 sequence: remaining, contains microbial sequence. For a 300 bp sequencing run, we use the following sequencing length configurations: read-1 sequence: 150 bp, the first 45 bp are barcode sequences, the last 75 bp are microbial sequences, those in the middle are adapter sequences; index-1 sequence: 8 bp; read-2 sequence: remaining, contains microbial sequence.

**Preprocessing of raw sequencing data**

We first group raw sequencing reads based on the 36864 barcodes. We exclude barcodes associated with too few reads, which collectively represent ~15% of the total reads. We also exclude those barcodes with a significantly more reads than other barcodes, which collectively represent ~5% of the total reads. For the remaining barcodes, we term the collection of microbial sequences associated with a single barcode as a single amplified genome (SAG). We use Trimmomatic (LEADING:25 TRAILING:3 SLIDINGWINDOW:4:20 MINLEN:30) to remove low quality reads and adapter sequences from each SAG for following analysis.

**Mock sample alignment, quality assessment, and coverage**

We use Bowtie2 (default parameters) to align reads from each SAG to the combined genome of the four reference genomes (RefSeq: GCF_002055965.1, GCF_004151095.1, GCF_001936035.1, GCF_002025145.1), which only reports the best hit of each read. We use SAMtools to check the number of reads that are aligned to each of the four genomes and to calculate the purity of each SAG. For each SAG with high purity (>0.95), we align reads from the SAG to the most aligned genome and to calculate the coverage of a SAG.

**Diversity of the human gut microbiome sample**s

For each of the seven samples, we temporarily ignore the barcode information and combine all reads from all SAGs from the sample. We use Kraken2 (default parameters) to classify reads from each *Microbe-seq* dataset and corresponding metagenomic dataset (standard Kraken database as of April 2019). For the analysis in Figure 3, we keep only the reads classified to a specific genus and use only this genus-level information for the comparison. However, analysis using all operational taxonomic units (OTUs) shows similar results (Table S2).

**Genome co-assembly of microbial species in the human gut microbiome**

We use SPAdes (--sc --careful) to *de novo* assemble genomes from the reads of each of the 21914 SAGs. We compute and compare signatures of these assembled genomes using sourmash (k-mer 51), which produces a matrix of estimated similarities between genomes. We use a hierarchical clustering method (method: complete, metric: Euclidean, criterion: 'inconsistent', and threshold: 0.95) to group SAGs into bins. We use this set of parameters because it groups bins conservatively, to minimize the improper grouping SAGs of different species together. We use all of the reads within each new bin to co-assemble a tentative genome, compare tentative genome similarities, and cluster the bins. We iterate this process until more than 10% of the assembled genomes have more than 10% contamination, estimated by CheckM (default parameters). Through four rounds, we group the 21914 SAGs into 364 bins.

We split bins that each contain SAGs from multiple species. Within each of the 364 bins, we align reads from each SAG to the *de novo* co-assembled tentative genome from the bin using bowtie2 (default parameters). For each contig in the tentative genome with more than 1000 bp, we construct a vector of each contig with the number of reads aligned to the contigs from each SAG. We use a hierarchical clustering method (method: ward) to group vectors of contigs into two groups. Although there might be more than 2 species in the bin, we apply binary clustering to each bin, as well as to each of the new sub-bins, to minimize supervision. For each SAG, if >95% aligned reads are aligned to one of the two groups of contigs, it is designated as a SAG associated with that group of contigs. We assign other SAGs to be a mixture of multiple species and exclude them from further analysis. We iterate this binary splitting process until we exclude more than 60% of the SAGs in the current bin, or both of the resulting new bins have fewer than than 10 SAGs, or the change between the resulting new bin and the current bin is less than three SAGs. We obtain 400 bins; SAGs within each bin are expected to be from the same species, with minimal level of contamination.

To combine bins of the same species for genome assembly, we use fastANI (default parameters) to calculate average nucleotide identity (ANI) between all pairs of these 400 genomes. Applying the commonly-used ANI > 95% threshold, above which two genomes are considered to represent the same species, we generate 234 new species-level bins. We *de novo* assemble reads from all SAGs within each of these 234 bins and remove contigs shorter than 500 bp. To further remove contigs that may originate from other species, within each genome, we fit a normal distribution with the coverage of contigs on a log scale and remove those contigs with coverages that are less than two standard deviations from the mean of the distribution.

Among these 234 genomes, 76 genomes are of high-quality (>90% completeness and <5% contamination) or medium-quality (>50% completeness and <10% contamination), as assessed by CheckM (default parameters). We use fastANI (default parameters) to compare the genomes of these 76 bins to all microbial genomes (RefSeq as of September 2019), and to the published collection of more than 1300 cultured-isolate whole genomes (Poyet et al., 2019). We identify the closest corresponding species-level genomes with ANI > 95% in both databases. The closest genomes in RefSeq to species *Alistipes onderdonkii*, *Bacteroides fragilis*, and *Bacteroides ovatus* are cultured isolate whole genomes from the same donor, reported earlier (Jiang et al., 2019); we exclude these three genome pairs from ANI and shared genome fraction analysis (Figure S3). We use BLASTn (default parameters) to compare overlapping sequences between genome pairs.

The names of the species-level genomes in RefSeq are not always labeled consistently; for example, we have four species that are named as *Blautia obeum* in RefSeq, though their ANI values are less than 95%. We use both GTDB-Tk (reference data version r89) and comparison to RefSeq genomes (September 2019) to assign taxonomies to all species. In the main text, we use taxonomies classified with GTDB-Tk and remove sub-genus names, such as '_A', for clarity.

**Phylogeny analysis of genomes**

To construct the phylogeny of the 76 species with high-quality or medium-quality genomes, we extract amino acid sequences of six ribosomal proteins (Ribosomal_L1, Ribosomal_L2, Ribosomal_L3, Ribosomal_L4, Ribosomal_L5, and Ribosomal_L6), concatenate and align them with Anvi'o. We construct a maximum-likelihood tree with RAxML (standard LG model, 100 rapid bootstrapping). We use iTOL to visualize and annotate the resulting dendrograms.

**Differentiate strains of the same species**

In the main text, the uncertainty in similarity of the bases at shared SNP locations in each pair of SAGs refers to the standard deviation of the normal approximation of the binomial distribution: uncertainty = sqrt[$p$(1-$p$)/n], where $p$ is the probability of the event and $n$ is the number of events. In the case of *Bacteroides vulgatus*, $n$=80 and the uncertainty is less than 6%.

Within each of the 30 bins of species with more than 100 SAGs, we align each SAG to the assembled genome from the bin.We use bcftools (mpileup, filters: snps and %QUAL>30) to identify high-quality single-nucleotide polymorphism (SNP) mutations. We apply the following criteria to process SNPs and SAGs, and perform these steps in order: a SAG with less than 2 reads aligned to a SNP is treated as unknown/unaligned at this location; a SAG with less than 99% reads being the same at a SNP is treated as unknown/unaligned at this location; a SNP with less than 5% of SAGs aligned to the location is removed; a SNP with less than two SAGs being the reference allele or less than two SAGs being the mutation allele is removed; a SNP with less than 1% SAGs being the reference allele or less than 1% SAGs being the mutation allele is removed; a SAG that is unknown at more than 99% of the kept SNP locations is removed; a SAG that is known at less than 10 of the kept SNP locations is removed.

We identify thousands of SNP locations and remove up to 6% of SAGs. We construct a SNP vector to represent the base identity sequence of each SAG at each SNP location. To identify the number of strains of the species in our samples, we build a dendrogram of SAGs with hierarchical clustering (method: 'ward') using the SNP vectors of all SAGs. Although the number of clusters is not obvious from the dendrogram, we obtain a sequence of SAGs from the dendrogram; in this sequence, SAGs with similar SNP sequences are closer. We compare similarities of SNP vectors between SAGs at their shared SNP locations and construct a similarity heatmap with SAGs ordered in the same sequence as the corresponding dendrogram. We observe block-diagonal squares in the heatmap, which indicates that SAGs within each square are closer to each other than to SAGs in other squares. Using the block-diagonal squares in the heatmap, we determine the number of strains, though this number can be difficult to establish precise for species with relatively few SAGS (<200) and potentially more than two strains. For *Blautia obeum* it is unclear whether there are 3 or 4 strains; for *Parasutterella excrementihominis,* it is unclear if there are 2 or 3 strains. We apply UMAP (default parameters) to the SNP data to create dimensional-reduction plots (Figure S5).

To remove SAGs that have reads from microbes of multiple strains, we construct the consensus genotype of each strain by comparing the SNP vectors of SAGs of the same strain. If more than 90% of the values at a SNP location from all SAGs within the strain are the same, we use the value for this SNP in the consensus genotype for the strain; otherwise we drop this SNP location for this strain. We compare the SNP vector of each SAG to the consensus genotype of each strain and assign strains to those SAGs that match more than 95% locations at the consensus genotype of only one strain, which excludes fewer than about 1% of the SAGs from a species. We co-assemble strain-resolved genomes with reads from all SAGs in each of these assigned strains with SPAdes using default parameters.

**Horizontal gene transfer analysis**

To detect HGT events between two strains, we use BLASTn (-perc_identity 99.98) to identify sequences that occur in both of their genomes that are more than 5000 bp. This detects recent HGT, thought to have happened within the last millennium, based on known mutation rates. (Zhao et al., 2019)

**Host-phage association analysis**

To identify SAGs that are associated with both crAssphage and a bacterial cell, we use bowtie2 (default parameters) to align reads in each SAG to the crAssphage genome (Refseq: GCF_000922395.1). We designate SAGs with more than 5% reads aligned to the crAssphage genome as containing significant crAssphage reads; we keep these SAGs and align the non-crAssphage reads of these SAGs to each of the 76 high- or medium-quality genomes, as well as the combined genome of these 76 genomes. We define purity of these SAGs as the maximum number of aligned reads to individual genomes divided by the number of aligned reads to the combined genome. We identify SAGs with more than 50% of reads aligned to one of these 76 genomes, and with purity of more than 95%. We designate the species of the SAG as the species of the most aligned genome. We count the number of SAGs assigned to each species and perform the "one species versus remaining species" one-sided Fisher's exact test. Among the SAGs with both bacterial reads and crAssphage reads, 14 are associated with *Bacteroides vulgatus*. We identify the strains of these 14 SAGs by comparing the SNP vectors of these SAGs to the consensus genotypes of each strain.


Klein, Allon M., Mazutis, L., Akartuna, I., Tallapragada, N., Veres, A., Li, V., Peshkin, L., Weitz, David A., and Kirschner, Marc W. (2015). Droplet Barcoding for Single-Cell Transcriptomics Applied to Embryonic Stem Cells. Cell *161*, 1187-1201.

McDonald, J.C., Duffy, D.C., Anderson, J.R., Chiu, D.T., Wu, H., Schueller, O.J.A., and Whitesides, G.M. (2000). Fabrication of microfluidic systems in poly(dimethylsiloxane). ELECTROPHORESIS *21*, 27-40.

Zilionis, R., Nainys, J., Veres, A., Savova, V., Zemmour, D., Klein, A.M., and Mazutis, L. (2017). Single-cell barcoding and sequencing using droplet microfluidics. Nature protocols *12*, 44.

Jiang, X., Hall, A.B., Arthur, T.D., Plichta, D.R., Covington, C.T., Poyet, M., Crothers, J., Moses, P.L., Tolonen, A.C., Vlamakis, H., *et al.* (2019). Invertible promoters mediate bacterial phase variation, antibiotic resistance, and host adaptation in the gut. Science *363*, 181-187.

Poyet, M., Groussin, M., Gibbons, S.M., Avila-Pacheco, J., Jiang, X., Kearney, S.M., Perrotta, A.R., Berdy, B., Zhao, S., Lieberman, T.D., *et al.* (2019). A library of human gut bacterial isolates paired with longitudinal multiomics data enables mechanistic microbiome research. Nature Medicine *25*, 1442-1452.

Zhao, S., Lieberman, T.D., Poyet, M., Kauffman, K.M., Gibbons, S.M., Groussin, M., Xavier, R.J., and Alm, E.J. (2019). Adaptive Evolution within Gut Microbiomes of Healthy People. Cell Host & Microbe *25*, 656-667.e658.
