## Supplementary material for "*Microbe-seq*: high-throughput, single-microbe genomics with strain resolution, applied to a human gut microbiome": microbeseq_supfig

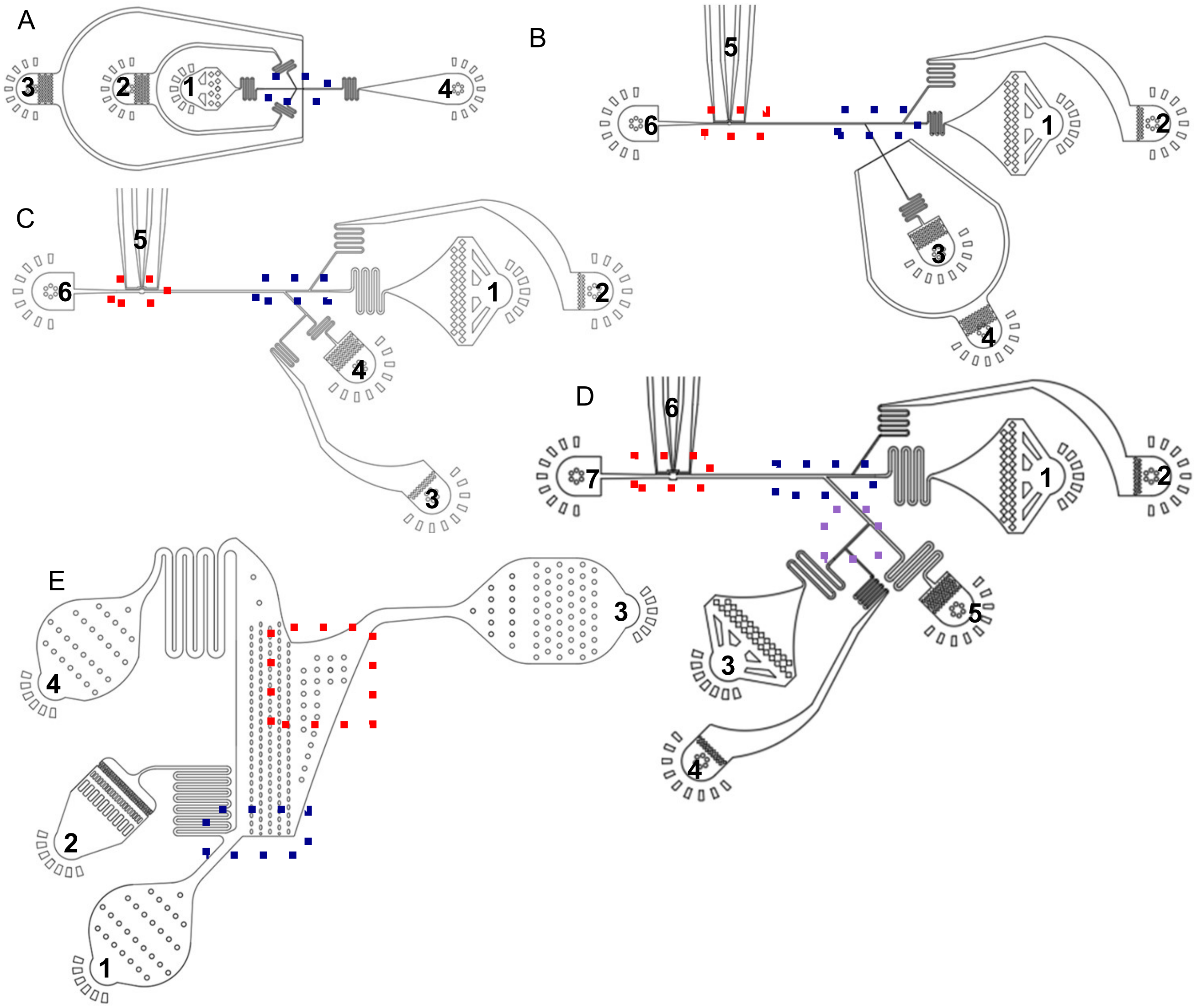


Figure S1. Designs of microfluidic devices for *Microbe-seq* platform, Related to Figure 1
(A) Design of the device to isolate microbes. Labels 1 to 4 are microbial suspension inlet, lysis reagent inlet, oil inlet, and collection outlet, respectively.
(B) Design of the device to add multiple displacement amplification (MDA) reagents into droplets. Labels 1 to 6 are sample droplet inlet, oil inlet, MDA reagent inlet, oil inlet, electrodes, and collection outlet, respectively.
(C) Design of the device to add transposomes (Nextera) into droplets to fragment and tag sample DNA. Labels 1 to 6 are sample droplet inlet, oil inlet, tagmentation reagent inlet, oil inlet, electrodes, and collection outlet, respectively.
(D) Design of the device to add beads and polymerase chain reaction (PCR) reagents into droplets. Labels 1 to 7 are sample droplet inlet, oil inlet, bead inlet, PCR reagent inlet, oil inlet, electrodes, and collection outlet, respectively.
(E) Design of the device to remove merged droplets. Labels 1 to 4 are sample droplet inlet, oil inlet, collection outlet, and waste outlet, respectively.
Videos of regions highlighted in dashed rectangles are shown in supplement video S1.

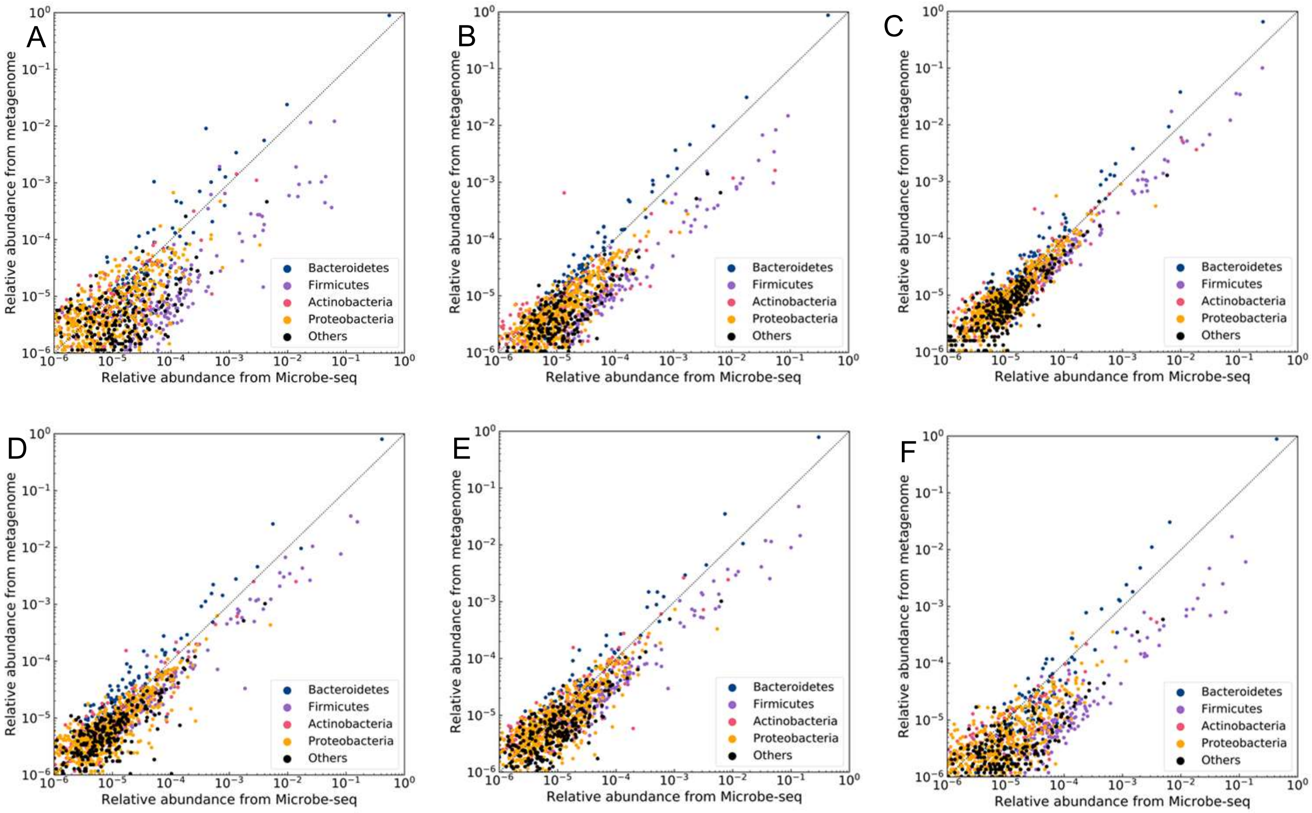


Figure S2. Microbial diversities in the human gut microbiome, Related to Figure 3.
Comparison of the relative abundance of genera captured by *Microbe-seq* and metagenome for samples 2 to 7 (A to F, respectively).

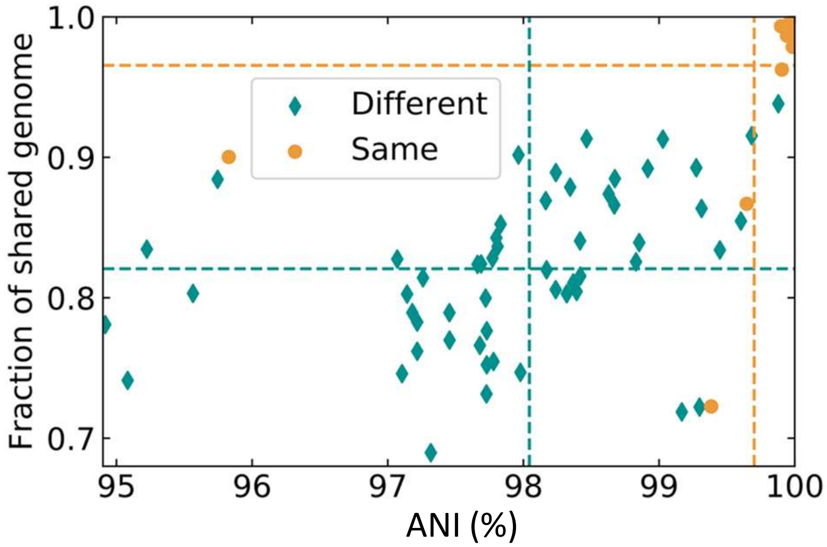


Figure S3. Similarity between co-assembled genomes and genomes of the same species, Related to Figure 5.

The similarity between co-assembled genomes and corresponding closest genomes (assessed by ANI) of the same species from different subjects (RefSeq) and the same subject. The fraction of shared genome is the fraction of genome sequence assembled from *Microbe-seq* that can be aligned to corresponding genome of the same species. Genomes from the same subject mostly show much higher similarity compare to those from different subjects.

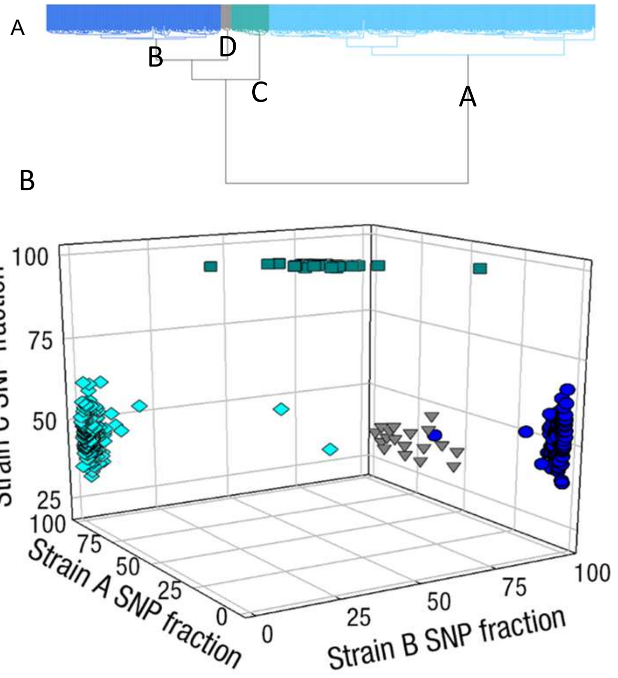


Figure S4 Strains of *Bacteroides vulgatus,* Related to Figure 5.
(A) Clustering of *Bacteroides vulgatus* SAGs into four strains. The dendrogram shows hierarchical clustering of SAGs based on their sequences at single nucleotide polymorphism (SNP) positions across the genome.

(B) Fraction of SNPs of each SAG that have the same base at the corresponding location in the three most populous clusters consensus genotypes.

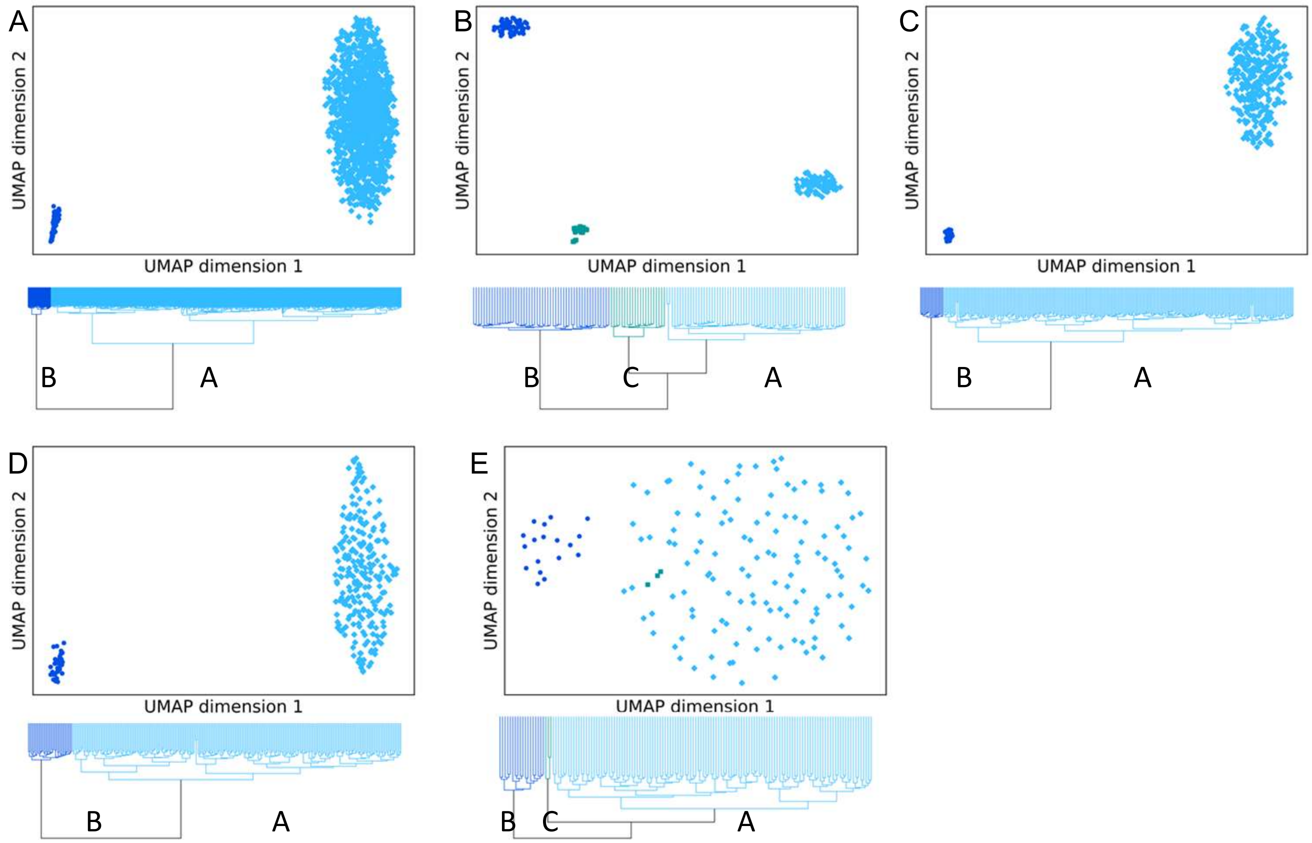


Figure S5. Strains of *Agathobacter faecis* (A), *Blautia_A obeum* (B), *Erysipelatoclostridium sp.* (C), *Faecalicatena faecis* (D), *and Parasutterella excrementihominis* (E), Related to Figure 5.

Each dendrogram shows hierarchical clustering of SAGs based on their sequences at single nucleotide polymorphism (SNP) positions across the genome. SNP sequences of these SAGs are visualized with dimension reduction (UMAP), SAGs in different groups from hierarchical clustering are in different colors.
